# GC-AG Introns Features in Long Non-coding and Protein-Coding Genes Suggest Their Role in Gene Expression Regulation

**DOI:** 10.1101/683938

**Authors:** Monah Abou Alezz, Ludovica Celli, Giulia Belotti, Antonella Lisa, Silvia Bione

## Abstract

Long non-coding RNAs (lncRNAs) are recognized as an important class of regulatory molecules involved in a variety of biological functions. However, the regulatory mechanisms of long non-coding genes expression are still poorly understood. The characterization of the genomic features of lncRNAs is crucial to get insight into their function. In this study, we exploited recent annotations by GENCODE to characterize the genomic and splicing features of long non-coding genes in comparison with protein-coding ones, both in human and mouse. Our analysis highlighted differences between the two classes of genes in terms of their gene architecture. Significant differences in the splice sites usage were observed between long non-coding and protein-coding genes (PCG). While the frequency of non-canonical GC-AG splice junctions represents about 0.8% of total splice sites in PCGs, we identified a significant enrichment of the GC-AG splice sites in long non-coding genes, both in human (3.0%) and mouse (1.9%). In addition, we found a positional bias of GC-AG splice sites being enriched in the first intron in both classes of genes. Moreover, a significant shorter length and weaker donor and acceptor sites were found comparing GC-AG introns to GT-AG introns. Genes containing at least one GC-AG intron were found conserved in many species, more prone to alternative splicing and a functional analysis pointed toward their enrichment in specific biological processes such as DNA repair. Our study shows for the first time that GC-AG introns are mainly associated with lncRNAs and are preferentially located in the first intron. Additionally, we discovered their regulatory potential indicating the existence of a new mechanism of non-coding and PCGs expression regulation.

## INTRODUCTION

The genomes of distantly related species house remarkably similar numbers of protein-coding genes (PCGs) prompting the notion that many aspects of complex organisms arise from non-coding regions (Liu et al., 2013; Fatica and Bozzoni, 2014). A large portion of mammalian genomes is transcribed to produce non-coding RNAs among which long non-coding RNAs (lncRNAs) are the most prevalent (Deveson et al., 2017). LncRNAs received growing attention as they emerged as an important regulatory layer of the transcriptome. They were described to be involved in transcriptional regulation, splicing, mRNA translation, chromatin modifications, and spatial conformation of chromosomes (Jandura and Krause, 2017; Mattick, 2018). Despite several studies reported their role in regulating the expression of other genes, how the transcription of lncRNAs is regulated remains less understood.

Similarly to PCGs, the majority of lncRNAs are transcribed by RNA Polymerase II and undergo the same RNA processing steps including capping, splicing, and polyadenylation. In comparison to PCGs, lncRNAs show lower levels of expression and higher tissue-specificity. The transcription of lncRNAs was mainly studied in relationship to those of nearby PCGs. In many cases, lncRNAs were reported to be co-expressed and co-regulated with their neighbor PCGs especially when divergently transcribed from bidirectional promoters (Sigova et al., 2013; Uesaka et al., 2014). In some cases, the direct involvement of lncRNAs in the transcription regulation of neighbor PCGs was demonstrated: in the work of Luo et al. (2016), the correlation between the expression of some lncRNAs and of the neighbor PCGs was experimentally demonstrated and estimated to account for 75% of total lncRNAs. As an example, the lncRNA EVX1-AS (EVX1-antisense RNA) was reported to promote the transcription of the EVX1 (even-skipped homeobox 1) gene during mesodermal differentiation by modifying chromatin accessibility (Luo et al., 2016). It was also reported that lncRNA transcription itself, rather than the RNA transcript, exerts regulatory effects on neighboring genes (Long et al., 2017). For example, the silencing of Igf2r (insulin like growth factor 2 receptor) gene expression was demonstrated to be due to the transcription of the lncRNAs Airn (antisense of Igf2r non-protein coding RNA) that interfere with RNA polymerase II recruitment (Latos et al., 2012). Similarly, Anderson et al. (2016) reported that the lack of transcription of the lncRNA Hand2os1 (Hand2, opposite strand 1), but not the knockdown of its mature transcript, abolished the expression of the Hand2 (heart and neural crest derivatives expressed 2) gene leading to embryonic lethality in mice. Taken together, this evidence points toward the tight regulation of lncRNAs expression and the importance of lncRNAs transcription in regulating PCGs.

Splicing represents a main mechanism of post-transcriptional regulation of gene expression (Papasaikas and Valcárcel, 2016). It is not only involved in the maturation of pre-mRNAs, but can also influence the subcellular localization of mature transcripts and increase transcriptional rates by several folds (Fong and Zhou, 2001). The majority of lncRNAs are processed by the splicing machinery and they can undergo alternative splicing showing a complexity in their gene expression regulation as mRNAs. Despite lncRNAs are less conserved than PCGs due to the absence of constraints on coding sequences (Hezroni et al., 2015), they exhibit conservation and selective constraints at their exon–intron structures and splicing regulatory elements (Schüler et al., 2014; Nitsche et al., 2015; Chernikova et al., 2016). Thus, the recognition of lncRNAs intron boundaries and the correct splicing of their introns is a crucial step for their functional maturation (Ponjavic et al., 2007; Nitsche and Stadler, 2017). Initial studies reported that lncRNAs show an overall splicing inefficiency compared with PCGs (Derrien et al., 2012; Tilgner et al., 2012; Melé et al., 2017). The inefficiency in lncRNAs splicing was mildly correlated to weak U2AF65 binding to 3′splice site (ss), in addition to the 5′ss strength and a lower thymidine content in the polypyrimidine tract of lncRNA introns (Melé et al., 2017; Krchòáková et al., 2019). Nevertheless, efficient splicing was observed among lncRNAs with specific functions (Melé et al., 2017). As well as for transcription, lncRNAs splicing can also affect the transcription of neighboring PCGs. In the study of Engreitz et al. (2016), it was demonstrated that the first 5′splice site of the mouse lncRNA Blustr has a critical impact on its ability to regulate the upstream PCG Sfmbt2 (Scm-like with four mbt domains 2). Thus, a better understanding of the mechanisms regulating lncRNAs splicing could contribute to understand their regulation and impact on PCGs transcription.

In this study, we took advantage of lncRNA annotations provided by the GENCODE project (Frankish et al., 2019) to characterize the genomic and splicing features of human and mouse lncRNAs in comparison to PCGs. At the genomic level, our analysis revealed differences in gene architecture between lncRNAs and PCGs, mainly in genic regions involved in gene expression regulation. The characterization of splicing features revealed a significant enrichment of GC-AG splice junctions in lncRNAs of human and mouse. Moreover, the GC-AG introns were preferentially found located in the first intron in both lncRNAs and mRNAs of both species. Based on the evidence that the frequency of 5′ss-GC was reported to increase with organisms complexity (Sheth et al., 2006) and that an accumulation of 5′ss-GC was previously described in mammals (Churbanov et al., 2008), we hypothesized that GC-AG introns may represent new key regulatory elements. Further analyses demonstrated that GC-AG introns differ from GT-AG introns in terms of length and donor and acceptor splice sites strength especially in lncRNAs. Interestingly, GC-AG introns appeared more prone to alternative splicing in both lncRNAs and mRNAs and in particular in alternative donor splice sites. In addition, GC-AG introns in PCGs appeared highly conserved and significantly enriched in specific biological processes such as DNA repair and neurogenesis. Taken together, our results highlighted unique features of GC-AG introns thus supporting their role as specific transcription regulators.

## MATERIALS AND METHODS

### Data collection

The lists of lncRNAs and PCGs were downloaded from the GENCODE website (https://www.gencodegenes.org/). Data from the release v27 were used for human genes annotated on the genome sequence GRCh38 (gencode.v27.long_noncoding_RNAs.gtf.gz; gencode.v27.basic.annotation.gtf.gz). Data from the release M16 were used for mouse genes annotated on the genome sequence GRCm38 (gencode.vM16.long_noncoding_RNAs.gtf.gz; gencode.vM16.basic.annotation.gtf.gz). PCGs were selected from the basic annotation when both gene and transcript were indicated as “protein_coding”. The total number of genes, transcripts and exons considered in both species are reported in Supplementary Table S1.

An independent validation of the results from GENCODE was obtained by collecting human lncRNAs annotations data from 6 different databases: the FANTOM5 database (Fantom CAT genes (http://fantom.gsc.riken.jp/cat/); FANTOM_CAT.lv3_robust.only_lncRNA.gtf) (Hon et al., 2017), the NONCODE v.5 database (http://noncode.org/datadownload/NONCODEv5_human_hg38_lncRNA.gtf.gz) (Fang et al., 2018), the BIGTranscriptome database release 2016 lncRNA catalog (http://big.hanyang.ac.kr/UCSC/RNA-seq/hg19/CAFE/GTFs/BIGTranscriptome/BIGTranscriptome_lncRNA_catalog.gtf) (You et al., 2017), the LncBook database (http://bigd.big.ac.cn/lncbook/index) (Ma et al., 2019), the MiTranscriptome database (http://mitranscriptome.org/download/mitranscriptome.gtf.tar.gz) (Iyer et al., 2015), and the LNCipedia database version 5.2 (https://lncipedia.org/downloads/lncipedia_5_2/full-database/lncipedia_5_2_hg38.gtf) (Volders et al., 2013). A validation of results obtained from the mouse genome was performed using lncRNAs annotations from the NONCODEv5 database (http://noncode.org/datadownload/NONCODEv5_mouse_mm10_lncRNA.gtf.gz).

The lists of lncRNAs and PCGs of Drosophila melanogaster and Caenorhabditis elegans were downloaded from the BioMart data mining tool (Smedley et al., 2015) in the Ensembl genome database (release 91).

### Conservation Analysis

To evaluate the conservation of genes containing GC-AG introns, we downloaded the list of orthologous genes in the human (GRCh38.p10) and mouse genomes (GRCm38.p5) from the Ensembl genome database (release 91) by using multi-species comparison in the BioMart data mining tool (Smedley et al., 2015). Multi-species conservation of 5′splice sites was assessed manually by aligning the sequences of corresponding introns in different organisms using the UCSC genome browser as data source (Kent et al., 2002). Species considered in this analysis were: human, chimp, macaque, mouse, rat, dog, cow, pig, chicken, fugu, and zebrafish.

### Introns Analysis

Intron sequences were retrieved using the Table Browser tool from UCSC using human GRCh38 and mouse GRCm38 genome sequences (Karolchik et al., 2004). We excluded from the analysis all single-exon genes as they are not subjected to splicing: this resulted in a total of 56582 lncRNAs and 525149 PCGs introns in human and 29611 lncRNAs and 393788 PCGs introns in mouse.

The scores of splice junctions were calculated using the MaxEntScan web tool (Yeo and Burge, 2004), a program for predicting the strength of the splicing sequences based on the maximum entropy model. In particular, MaxEntScan::score5ss scores the donor splice site from a sequence motif of 9 nucleotides covering bases −3 to +6 and accounts for non-adjacent as well as adjacent dependencies between positions. MaxEntScan::score3ss scores the acceptor splice site from a sequence motif of 23 nucleotide covering bases −20 to +3. We evaluated the strength of 5′ and 3′ splice sites of human and mouse introns using the Weight Matrix Model as provided by the MaxEntScan tool. The evaluation of the polypyrimidine tract strength was performed using the “branchpointer” R package version 1.10.0 (Signal et al., 2018). The package predicted polypyrimidine tracts in query regions located at −18 to −44 nucleotides from the 3′ splice sites.

### Alternative Splicing Analysis

The assignment of alternative splicing events involving GC-AG and GT-AG introns was performed using the SUPPA2 tool (Trincado et al., 2018). Splicing events were extracted from the gtf files of lncRNA and PCGs annotations from the GENCODE database. The SUPPA2 tool classified alternative spliced events according to the following types: exon skipping, intron retention, mutually exclusive exons, alternative 5′ss, alternative 3′ss, alternative first exons and alternative last exons. Custom R scripts were used to extract introns involved in each type of alternative splicing event and to evaluate alternative last exons. Polyadenylation signals (PAS) were extracted according to the 16 PAS reported in the paper of Beaudoing et al. (2000) in a bin of 40 nucleotides at the end of each last exon.

### Expression Analysis

RNA-Seq data of healthy individuals were obtained from the Genotype-Tissue Expression (GTEx) version 8 data set (https://www.ncbi.nlm.nih.gov/projects/gap/cgi-bin/study.cgi?study_id=phs000424.v8.p2; phs000424.v8.p2.c1, July 18, 2019) (GTEx Consortium, 2015) and downloaded using dbGaP web site (approved protocol #23403). Data were collected from 10 different tissues (anterior cingulate cortex, amygdala, cerebellum, heart left ventricle, kidney cortex, lung, liver, spleen, skin, and testis) of male individuals using 8 samples per tissue for a total of 80 samples. Quality control analyses on the raw sequence data were performed using the FastQC tool (http://www.bioinformatics.babraham.ac.uk/projects/fastqc/). Reads in FASTQ format passing quality control were quantified with the transcripts per million method implemented in the Salmon software (version 1.2.0) (Patro et al., 2017) using default parameters and the human hg38 reference transcriptome from GENCODE v27. The transcripts quantifications were then imported into R and summarized using custom scripts. The unexpressed lncRNA transcripts with TPM < 0.1 and proteincoding transcripts with TPM < 0.5 were filtered out in subsequent analyses.

### Statistical Analysis

Data analyses and descriptive statistics were performed using RStudio version 1.1.456 (http://www.rstudio.com/). The Wilcoxon rank-sum test was applied to compare distributions and the Chi-square test was applied to compare groups. Correlation analysis was performed by estimating the Spearman correlation coefficient (r). For all statistical tests, a p-value < 0.05 was considered as significant.

### Functional Enrichment Analysis

Gene list functional enrichment analyses were performed using the DAVID (Database for Annotation, Visualization and Integrated Discovery; version 6.8) tool (Huang et al., 2009) and the PANTHER (“Protein ANalysis THrough Evolutionary Relationships”; release 20181113) overrepresentation test (Mi et al., 2019) implemented in the Gene Ontology (GO) website (Ashburner et al., 2000; The Gene Ontology Consortium, 2019). The lists of PCGs containing a GC-AG intron from both human (n = 1934) and mouse (n = 1669) were subjected to an enrichment analysis on GO Biological Process terms and filtered applying a statistical significance threshold of 0.05 based on the multiple testing corrected p-values [i.e., Benjamini adjusted p-value in DAVID or false-discovery rates (FDR) in PANTHER].

### Availability of Data and Materials

This study was based on genomic data of lncRNAs and PCGs provided by the GENCODE Project. In particular, data of lncRNAs and PCGs were downloaded from the GENCODE web-pages for human (https://www.gencodegenes.org/human/release_27.html) and mouse (https://www.gencodegenes.org/mouse/release_M16.html) as gtf files. We only analyzed anonymized samples for which the corresponding donor consent information was available in the GTEx dataset (dbGaP:phs000424.v8.p2) at the time of the analysis. Samples were downloaded from the dbGap database (https://www.ncbi.nlm.nih.gov/gap) according to the specified guidelines. All of the samples we analyzed were approved for General Research Use (GRU) and thus have no further limitations outside of those in the NIH model Data Use Certification Agreement.

The datasets supporting the conclusions of this article (i.e., introns data) are available in the GitHub repository (https://github.com/laBione/introns_data)

## RESULTS

### Long non-coding and protein-coding genes showed differences in their gene structure

As genomic organization and gene structure may affect gene expression regulation, we characterized the genomic features of human and mouse lncRNAs in comparison with PCGs. Our analysis, based on GENCODE human release 27 (15778 lncRNAs and 19836 PCGs) and mouse release M16 (12374 lncRNAs and 21963 PCGs), considered an increased number of genes with respect to previous studies (Cabili et al., 2011; Derrien et al., 2012). The total number of genes, transcripts and exons are reported in Supplementary Table S1.

The genomic organization of lncRNAs and PCGs appeared highly similar in both species. Human and mouse lncRNAs appeared equally transcribed from the forward and the reverse strand as PCGs (Supplementary Table S2) and almost homogeneously interspersed along chromosomes. Gene density resulted highly variable among chromosomes for lncRNAs and PCGs in both species (Supplementary Table S3 and Supplementary Figure S1).

The genome coverage of long non-coding genes was found remarkably lower with respect to that of protein-coding ones. Indeed, long non-coding genes accounted for 12.5% of the human genome while 43.4% is occupied by PCGs (Chi-square test = 730.4, 1 df, p-value < 2.2 × 10–16). The reduced genome coverage was not entirely due to the smaller number of lncRNAs, as they account for about 80% of protein-coding ones, but it appeared to be due to the lncRNAs length, that resulted significantly lower than that of PCGs. Human lncRNAs resulted, on average, almost three times shorter than protein-coding ones with an average length of about 24 kb versus 68 kb, respectively (Wilcoxon test p-value < 2.2 × 10–16) (Figure 1A and Supplementary Table S4). Similarly, the genome coverage of mouse lncRNAs was lower (6.8%) than that of PCGs (39.2%) (Chi-square test = 802.5, 1 df, p-value < 2.2 × 10–16). The lower genome coverage was not only due to the smaller number of lncRNAs (accounting for 56% of PCGs) but also to their gene length that, as in human, resulted significantly shorter than that of PCGs with an average length of about 15 kb versus 49 kb, respectively (Wilcoxon test p-value < 2.2 × 10–16) (Figure 1B and Supplementary Table S4).

**Figure 1.**
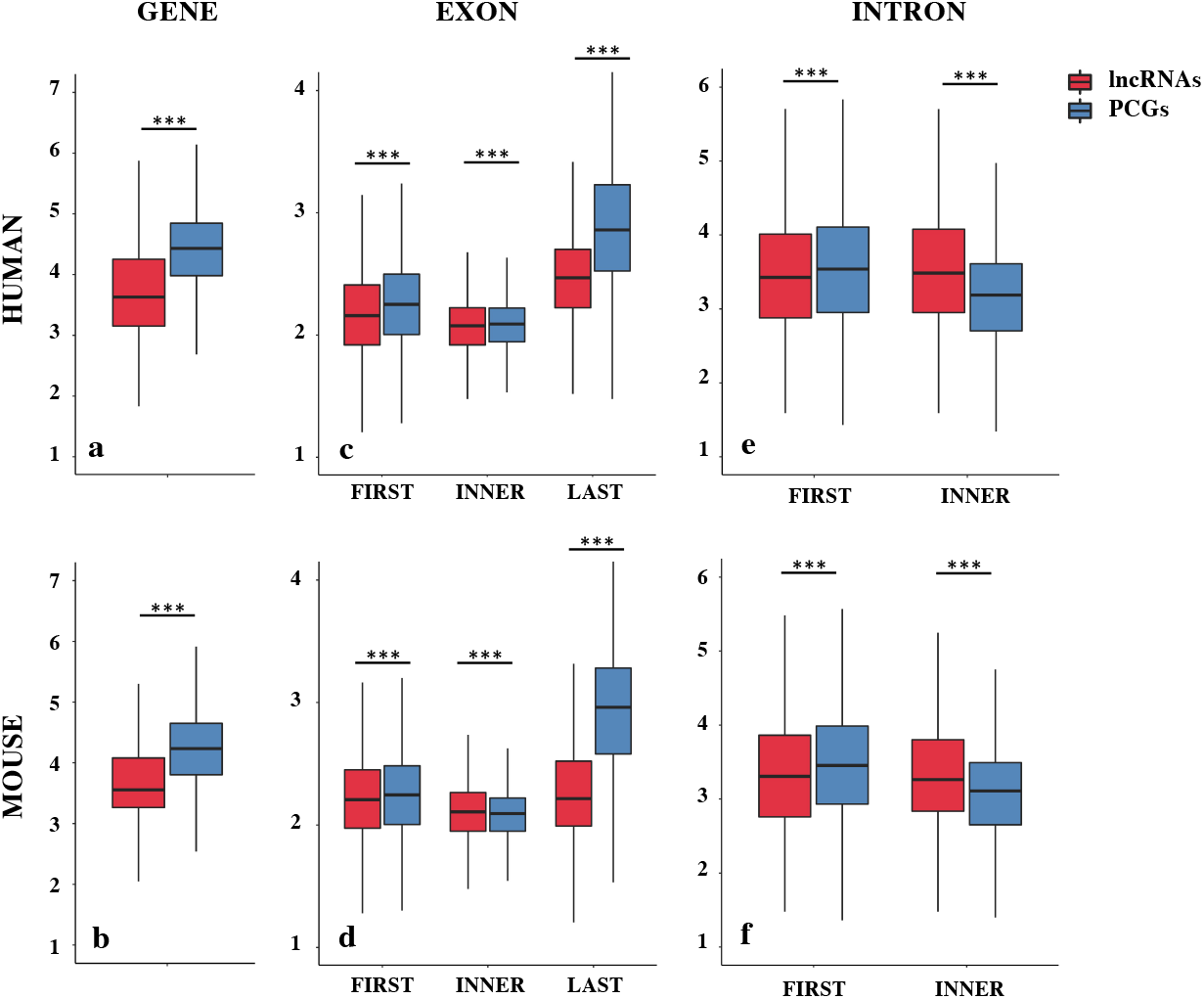
Gene structure features of long non-coding and protein-coding genes (PCGs) in human and mouse. Boxplots showing the gene length (A,B), exon length (C,D), and intron length (E,F) in human and mouse, respectively. Data were presented as log10 of length in base pairs (bp). Exons were classified as first, inner or last and introns were classified as first or inner. ***p < 0.001.

The shorter length of lncRNAs was attributable to the lower number of exons composing them (Supplementary Table S5). In human, more than 70% of lncRNA transcripts had 3 exons or less, compared with 16% of protein-coding transcripts bearing the same characteristics (Chi-squared test = 24407.0, 1 df, p-value < 2.2 × 10–16). A large proportion of lncRNA transcripts was composed of 2 exons (34%) as previously reported (Derrien et al., 2012) and 14% are single-exon genes. In mouse, more than 75% of lncRNAs had 3 exons or less versus 23% in protein-coding transcripts (Chi-squared test = 14613.7, 1 df, p-value < 2.2 × 10–16) and 24% of lncRNAs were single-exon genes versus 6.4% in protein-coding ones. Also in the mouse genome, an enrichment of 2-exons transcripts (30%) was observed (Supplementary Table S5). These results were confirmed using the FANTOM5 collection of human lncRNAs, an independent source for the annotation of lncRNAs (Hon et al., 2017) in which we observed the same trend for lncRNAs length (mean = 28.2 kb, SEM = 458.4 bp) and lower number of exons (less than 3 exons: 56%).

A deep characterization of exons and introns length allowed us to appreciate differences between lncRNAs and PCGs. Conversely to what was previously reported in Derrien et al. (2012), our data revealed that first exons and especially last exons in lncRNAs are significantly shorter in both species (Supplementary Table S6 and Figures 1C,D). LncRNA introns were found longer than PCGs ones when they are inner introns; instead, they resulted slightly shorter when they are first introns (Supplementary Table S6 and Figures 1E,F).

Although it is possible that these differences were due to an incomplete annotation of lncRNAs, it is nevertheless interesting to note that the reduction in size affects those portions of the gene mainly involved in gene expression regulation.

### Different Assortment of Splicing Junctions Consensus in Long Non-coding RNAs

As splicing is a main determinant of post-transcriptional gene expression regulation, we characterized the splicing features of lncRNA introns in comparison with those of protein-coding ones.

The splice junctions sequence analysis highlighted differences between lncRNAs and PCGs consensus sequences (Table 1). The GC-AG splice junctions appeared strongly enriched in lncRNAs in which they represent 3.0% of the total splice junctions, thus almost four times more than in PCGs (0.8%) (Chi-square test = 2289.4, 1 df, p-value < 2.2 × 10–16). The same enrichment was found in mouse, in which GC-AG splicing junctions were more than the double with respect to protein-coding ones (lncRNAs: 1.9%, pc 0.8%) (Chi-square test = 380.2, 1 df, p-value < 2.2 × 10–16).

**Table 1.** Number of different splice junctions consensus.

|  | **Human** |  |  |  |
| --- | --- | --- | --- | --- |
|  | **#lncRNAs** | **%** | **#PCGs** | **%** |
| **GT-AG** | 54667 | 96.6 | 517730 | 98.6 |
| **GC-AG** | 1683 | 3.0 | 4351 | 0.8 |
| **AT-AC** | 9 | 0.0 | 583 | 0.1 |
| **Others** | 223 | 0.4 | 2485 | 0.5 |
| **Total** | 56582 |  | 525149 |  |
|  | **Mouse** |  |  |  |
|  | **#lncRNAs** | **%** | **#PCGs** | **%** |
| **GT-AG** | 28586 | 96.5 | 388973 | 98.8 |
| **GC-AG** | 570 | 1.9 | 3217 | 0.8 |
| **AT-AC** | 6 | 0.0 | 363 | 0.1 |
| **Others** | 449 | 1.5 | 1235 | 0.3 |
| **Total** | 29611 |  | 393788 |  |

GC-AG introns showed a preferential location in the first intron of both lncRNAs and PCGs (Table 2). Indeed, in the human genome, their percentage resulted higher in the first intron (lncRNAs: 4.2%; PCGs: 1.2%) with respect to inner introns (lncRNAs: 2.1%; PCGs: 0.8%) and the same trend was observed in mouse (first: lncRNAs 2.4%, PCGs: 1.2%; inner: lncRNAs 0.4%, PCGs 0.8%). In all cases, differences were statistically significant (Chi-square tests = 204.7 and 120.9, 1 df, p-value < 2.2 × 10–16, respectively, for human lncRNAs and PCGs; Chi-square tests = 233.6 and 62.7, 1 df, p-value < 2.2 × 10–16 and < 2.4 × 10–15, respectively, for mouse lncRNAs and PCGs).

**Table 2.** Number of GC-AG introns in first or inner positions.

| **Human** |  |  |  |  |  |  |
| --- | --- | --- | --- | --- | --- | --- |
|  | **lncRNAs** |  |  | **PCGs** |  |  |
|  | **#** | **#GC-AG** | **%** | **#** | **#GC-AG** | **%** |
| **First** | 23997 | 1000 | 4.2 | 53776 | 665 | 1.2 |
| **Inner** | 32585 | 683 | 2.1 | 471373 | 3686 | 0.8 |
| **Total** | 56582 | 1683 | 3.0 | 525149 | 4351 | 0.8 |

| **Mouse** |  |  |  |  |  |  |
| --- | --- | --- | --- | --- | --- | --- |
|  | **lncRNAs** |  |  | **PCGs** |  |  |
|  | **#** | **#GC-AG** | **%** | **#** | **#GC-AG** | **%** |
| **First** | 13079 | 309 | 2.4 | 40990 | 472 | 1.2 |
| **Inner** | 16532 | 61 | 0.4 | 352798 | 2745 | 0.8 |
| **Total** | 29611 | 570 | 1.9 | 393788 | 3217 | 0.8 |

A validation of these results was obtained by investigating six alternative source of lncRNA annotations: (1) the FANTOM5 dataset (number of transcripts = 161340), (2) the NONCODE dataset (number of transcripts = 257020), (3) the BIGTranscriptome dataset (number of transcripts = 61018), (4) the LncBokk dataset (number of transcripts = 410630), (5) the MITranscriptome dataset (number of transcripts = 364544), and (6) the LNCipedia dataset (number of transcripts = 229818). In all datasets, the frequency of GC-AG splice junctions was found higher with respect to that in PCG introns together with their preferential location as the first intron (Supplementary Table S7). The prevalence of GC-AG introns among lncRNAs new datasets ranged from 2.3 to 3.5%, resulting in all cases significantly higher respect to PCGs (all comparisons p-values < 2.2 × 10–16). In all datasets, GC-AG introns showed their preferential localization in the first introns in which their prevalence in constantly the double with respect to inner introns (all comparisons p-values < 2.2 × 10–16). In mouse, data was replicated in the NONCODE dataset (Supplementary Table S8) in which both the enrichment of GC-AG splice junction (1.7% with respect to 0.8% in PCGs; p-values < 2.2 × 10–16) and their preferential location in the first intron (fist introns 2.3%, inner introns 1.1%; p-values < 2.2 × 10–16) was confirmed.

To evaluate the enrichment of GC-AG introns in lncRNAs during evolution, we analyzed the frequency of the different splice junctions in lower organisms as D. melanogaster and C. elegans. The ratio of GC-AG splice sites in lncRNAs of D. melanogaster was found significantly higher than in PCGs (GC-AG in lncRNAs: 1.7% of total splice junctions with respect to GC-AG in PCGs: 0.7%; Chi-square test = 57.0, 1 df, p-value = 4.3 × 10–14). In C. elegans, GC-AG splice junctions account for 2.0% of total splice junctions in lncRNAs thus confirming the enrichment with respect to the 0.6% in PCGs (Chi-square test = 12.7, 1 df, p-value = 3.5 × 10–4) (Supplementary Table S9). A preferential location of GC-AG splice sites in the first intron was also observed in lncRNA and PCGs of both D. melanogaster and C. elegans but due to their small number their statistical relevance could not be appreciated.

### Peculiar Features of GC-AG Introns in Long Non-coding and Protein-Coding Genes

The enrichment of GC-AG junctions in lncRNAs together with their preferential localization in first introns in both lncRNAs and PCGs suggested that they could play a particular role in gene expression regulation leading us to a deeper characterization of their features.

In human, GC-AG introns resulted shorter both in lncRNAs and PCGs and they showed the same trend whether they are first or inner introns (Supplementary Table S10). For GC-AG first introns, the average length resulted almost halved with respect to GT-AG first introns in both human lncRNAs and PCGs (lncRNAs: GC 6700 ±600 bp, GT 12923 ±201 bp, Wilcoxon tests p-value < 2.2 × 10–16; PCGs: GC 8999 ±648 bp, GT 15335 ±162 bp, Wilcoxon tests p-value < 2.2 × 10–16). Human GC-AG inner introns showed the same decrease in length, albeit to a lesser extent (lncRNAs: GC 8666 ±827 bp, GT 13995 ±194 bp, Wilcoxon tests p-value = 0.012; PCGs: GC 4165 ±197 bp, GT 5411 ±25 bp, Wilcoxon tests p-value = 6.3 × 10–10). In mouse, GC-AG introns appeared shorter but only when they are inner introns (lncRNAs: GC 5190 ±734 bp, GT 7523 ±148 bp, Wilcoxon tests p-value = 0.0302; PCGs: GC 3186 ±192 bp, GT 4437 ±27 bp, Wilcoxon tests p-value = 9.5 × 10–14). The shorter length of human GC-AG introns was also confirmed in the FANTOM5 dataset as both GC-AG first and inner introns of lncRNAs were significantly shorter than GT-AG ones (first intron: GC 8169 ±600 bp, GT 14516 ±137 bp, Wilcoxon tests p-value < 2.2 × 10–16; inner introns: GC 8648 ±544 bp, GT 15784 ±119 bp, Wilcoxon tests p-value < 2.2 × 10–16) (Supplementary Table S11).

To evaluate the splicing efficiency of GC-AG junctions, we computed their strength using the standard position weight-matrix (WM) model implemented in the MaxEntScan tool (Yeo and Burge, 2004), which assigns a computationally predicted score for 5′ and 3′ splice sites. Overall, the strength of 5′and 3′ss resulted lower in lncRNAs than in PCGs both in human and mouse (Figure 2, Supplementary Table S12, and Supplementary Figure S2) and it was presumably one of the causes of the previously reported inefficiency of lncRNAs splicing (Tilgner et al., 2012; Melé et al., 2017). Despite lower weight-matrix scores for 5′ss-GC were expected, due to their imperfect pairing with the U1 snRNA, 5′ss-GC scores of lncRNAs resulted strongly reduced with respect to 5′ss-GC of PCGs in both species (human: lncRNAs 5′ss-GC WM = 0.50, PCGs 5′ss-GC WM = 2.76, Wilcoxon test p-value < 2.2 × 10–16; mouse: lncRNAs 5′ss-GC WM = 1.63, PCGs 5′ss-GC WM = 3.38, Wilcoxon test p-value < 2.2 × 10–16). The reduced strength of lncRNAs 5′ss-GC appeared to be attributable almost exclusively to first intron junctions, whose scores resulted lower compared to those of inner introns, both in human and mouse (human: lncRNAs first intron 5′ss-GC WM = −0.93, inner intron 5′ss-GC WM = 2.60, Wilcoxon test p-value < 2.2 × 10–16; mouse: lncRNAs first intron 5′ss-GC WM = 0.78, inner intron 5′ss-GC WM = 2.65, Wilcoxon test p-value < 2.2 × 10–16).

**Figure 2.**
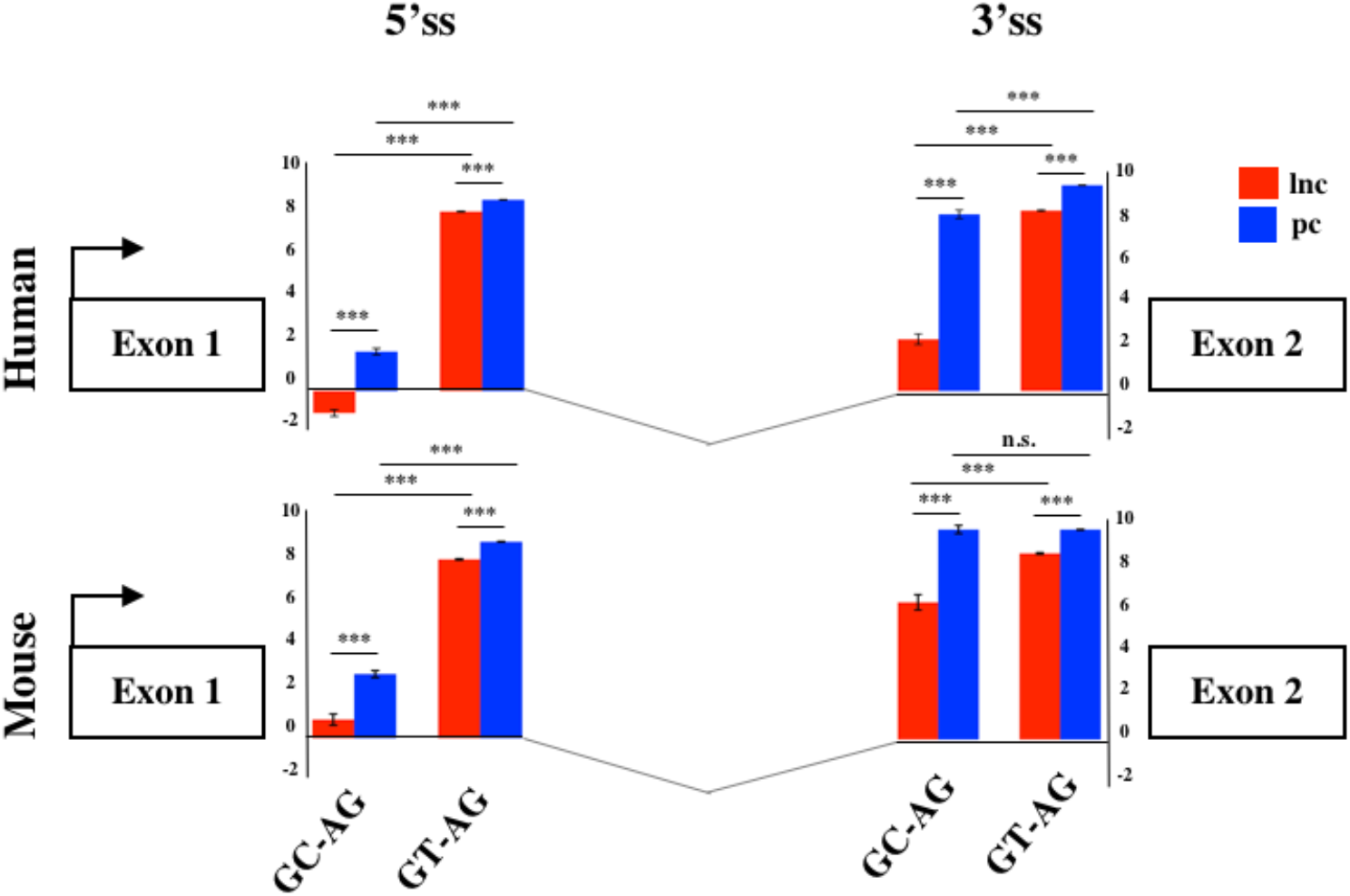
Splice junctions strengths of the first introns. Schematic representation of the average scores of 5′ and 3′ss strengths of long non-coding and PCGs in human and mouse. The strengths of 5′ and 3′ ss were calculated as weight matrix scores for GC-AG and GT-AG first introns. ***p < 0.001.

Despite owning the same consensus sequence, the 3′ss average weight-matrix scores for GC-AG introns appeared overall lower with respect to GT-AG acceptor sites and this appeared attributable to their shorter polypyrimidine tracts (PPT) (Supplementary Table S13). In human, the mean length of PPT of GC-introns resulted significantly shorter than GT ones in both lncRNAs and PCGs (lncRNAs: GT-introns PPT mean = 16 bp, GC-introns PPT mean = 12 bp, Wilcoxon tests p-value < 2.2 × 10– 16; PCGs: GT-introns PPT mean = 16 bp, GC-introns PPT mean = 15 bp, Wilcoxon tests p-value < 2.2 × 10–16). The same trend was observed in mouse for both gene classes (lncRNAs: GT-introns PPT mean = 15 bp, GC-introns PPT mean = 14 bp, Wilcoxon tests p-value = 0.001; PCGs: GT-introns PPT mean = 16 bp, GC-introns PPT mean = 15 bp, Wilcoxon tests p-value = 0.021). As it occurred for 5′ss, very weak 3′ss appeared preferentially located in the lncRNAs first intron in both human and mouse.

To test whether the 5′ss and 3′ss weight-matrix scores and the introns length showed any correlation, the Spearman test was applied (Supplementary Table S14). The strength of 5′ss and 3′ss was found positively correlated when located in the first intron of human lncRNAs (r = 0.58, p-value < 2.2 × 10–16) and PCGs (r = 0.22, p-value = 1 × 10–16). In mouse, the correlation was significant only in lncRNAs (lncRNAs: r = 0.51, p-value < 2.2 × 10–16; PCGs: : r = 0.04, p-value = 0.34). The strengths of both 5′ss and 3′ss were positively correlated to intron length and this correlation was found more pronounced in the first intron in both species.

Differently from what was reported for PCGs, in which weak donor sites appeared flanked by stronger consensus at the acceptor sites (Thanaraj and Clark, 2001; Kralovicova et al., 2011), our analysis demonstrated that lncRNAs contained a class of very weak introns, preferentially located as first.

### GC-AG Introns in Alternative Splicing and Polyadenylation Regulation

As the presence of a GC-AG intron was proposed to increase the level of alternative splicing (Churbanov et al., 2008), we compared the transcriptional diversity of both lncRNAs and PCGs owning at least one GC-AG intron with respect to the ones containing only GT-AG introns (Supplementary Table S15). In human, both long non-coding and protein-coding GC-AG-containing genes being transcribed in more than one isoform exceeded the number of GT-AG-containing genes [lncRNAs-GC n = 471 (38.5%) vs. lncRNAs-GT n = 3204 (28.9%), Chi-square test = 47.7, 1 df, p-value = 4.8 × 10–12; PCGs-GC n = 1642 (84.9%) vs. PCGs-GT n = 11469 (68.9%), Chi-square test = 212.5, 1 df, p-value < 2.2 × 10–16]. The same trend was confirmed in mouse (Supplementary Table S15), where long non-coding and protein-coding GC-AG-containing genes with more than one isoform resulted more abundant than their GT-AG counterpart [lncRNAs-GC n = 188 (39.7%) vs. lncRNAs-GT n = 2085 (25.3%), Chi-square test = 47.7, 1 df, p-value = 4.9 × 10–12; PCGs-GC n = 1117 (66.9%) vs. PCGs-GT n = 9463 (50.2%), Chi-square test = 170.6, 1 df, p-value < 2.2 × 10–16]. To evaluate if the increase of alternative splicing could be attributed to some particular splicing events, we used the SUPPA2 tool (Trincado et al., 2018) to perform a quantitative profiling of alternative splicing events involving GC-AG introns in comparison with GT-AG ones (Supplementary Table S16). The analysis revealed that human GC-AG introns were preferentially involved in the alternative 5′ss events in both lncRNAs and PCGs [lncRNAs: n = 150 (18.9%) of GC-AG introns, n = 3494 (9.7%) of GT-AG introns, Chi-square test = 54.9, 1 df, p-value = 1.2 × 10–13; PCGs: n = 389 (31.6%) of GC-AG introns, n = 10500 (10.1%) of GT-AG introns, Chi-square test = 415.6, 1 df, p-value < 2.2 × 10–16]. The same trend was also observed in mouse [lncRNAs: n = 41 (34.7%) of GC-AG introns, n = 1,000 (11.1%) of GT-AG introns, Chi-square test = 40.9, 1 df, p-value = 1.5 × 10–10; PCGs: n = 188 (32.9%) of GC-AG introns, n = 5902 (12.5%) of GT-AG introns, Chi-square test = 136.7, 1 df, p-value < 2.2 × 10–16].

As alternative polyadenylation regulation is a process directly linked to 5′ss recognition and splicing, we analyzed the variability of last exon (LE) defining the total number of alternative last exon for each gene. In this analysis, we assessed an overall enrichment of LE variability in PCGs respect to lncRNAs in both species (Supplementary Table S17A). Interestingly, we observed a significant increase of LE variability in GC-AG-containing genes compared to GT-AG ones in both gene classes. In human lncRNAs, 37.6% of GC-AG-containing genes had more than one alternative last exon compared to 27.7% of GT-AG-containing genes (Chi-square test = 52.4, 1 df, p-value = 4.5 × 10– 13). The same difference was established for human PCGs (80% of GC-AG genes with alternative last exon versus 64.1% of GT-AG genes, Chi-square test = 151.4, 1 df, p-value < 2.2 × 10–16). The same significant enrichment were confirmed in mouse (lncRNAs: 38.3% of GC-AG genes with alternative last exon versus 23.5% of GT-AG genes, Chi-square test = 52.4, 1 df, p-value = 4.5 × 10– 13; PCGs: 59.3% of GC-AG genes with alternative last exon versus 43.7% of GT-AG genes, Chisquare test = 151.4, 1 df, p-value < 2.2 × 10–16) (Supplementary Table S17B). Furthermore, the increased of LE variability in GC-AG-containing genes was strengthened by a higher mean of alternative last exons per gene respect to GT-AG-containing genes in human and mouse lncRNAs and PCGs (Supplementary Table S17C). As differences in polyadenylation regulation could result from the different assortment of polyadenylation signals (PAS), we analyzed the last 40 nucleotides of each last exon for their content in the 16 different PAS reported in the paper of Beaudoing et al. (2000). Our results highlighted a higher ratio of lncRNAs lacking any of the 16 PAS considered compared with PCGs in both species (human: lncRNAs PAS = 0 48.0% versus PCGs PAS = 0 29.7%, Chi-square test = 2200.3, 1 df, p-value < 2.2 × 10–16; mouse: lncRNAs PAS = 0 42.4% versus PCGs PAS = 0 20.3%, Chi-square test = 2370.9, 1 df, p-value < 2.2 × 10–16) (Supplementary Table S18A). Considering GC-AG- and GT-AG-containing genes separately, we observed that the higher ratio of PAS = 0 was more evident in GC-AG transcripts but the difference was statistically significant only in human (Supplementary Table S18B). Looking at the assortment of different PAS, we observed a preferential usage of non-canonical PAS in lncRNAs with respect to PCGs in both species (human: lncRNAs non-canonical PAS 65.4% versus PCGs non-canonical PAS 57.6%, Chi-square test = 346.1, 1 df, p-value < 2.2 × 10–16; mouse: lncRNAs non-canonical PAS 65.8% versus PCGs non-canonical PAS 56.7%, Chi-square test = 309.2, 1 df, p-value < 2.2 × 10–16) (Supplementary Table S18C). No differences between GC-AG- and GT-AG-containing genes were observed in the usage of different PAS (data not shown).

Our results highlighted differences in alternative splicing and polyadenylation sites and signals between lncRNAs and PCGs which appeared more evident in GC-AG-containing genes thus suggesting that this 5′ss could contribute to gene expression regulation.

### Impact of GC-AG Introns on Gene Expression Level

In order to evaluate a putative effect of the presence of a GC-AG intron on the expression level of the corresponding transcripts, we analyzed a panel of ten different human tissues (i.e., anterior cingulate cortex, amygdala, cerebellum, heart, kidney, liver, lung, skin, spleen, and testis) obtained from the GTEx project. For each tissue, raw RNA-seq data from eight samples were processed using the Salmon tool (Patro et al., 2017) which provide an accurate quantification of transcripts expression. Transcript per million (TPM) of each single transcript, were calculated in each tissue and expressed transcripts were defined based on a threshold of TPM > 0.1 for lncRNAs and of TPM > 0.5 for PCGs to account for highly different level of expression between the two classes of genes. The percentage of expressed transcripts and the mean TPM in each tissue were reported distinguishing between GC-AG- or GT-AG-intron containing transcripts and between transcripts containing a GC-AG intron in the first or inner position (Supplementary Table S19 and Supplementary Figure S4). In addition, we calculated the mean TPM of all tissues combined together in order to provide an overall estimation of expression data.

The mean TPM of GC-AG-containing transcripts appeared always lower with respect to GT-AG containing ones (with the exception of TPM values for lncRNAs in lung) and in the majority of the cases the difference resulted statistically significant. Combining all tissues together, the mean TPM of lncRNAs resulted significantly lower with respect to GT-AG-containing transcripts in both lncRNAs and PCGs (lncRNAs: 1.79 for GC-AG containing transcripts vs. 2.00 for GT-AG containing ones, Wilcoxon test p-value = 3.2 × 10–15; PCGs: 8.40 for GC-AG containing transcripts vs. 11.10 for GT-AG containing ones, Wilcoxon test p-value < 2.2 × 10–16) (Figure 3).

**Figure 3.**
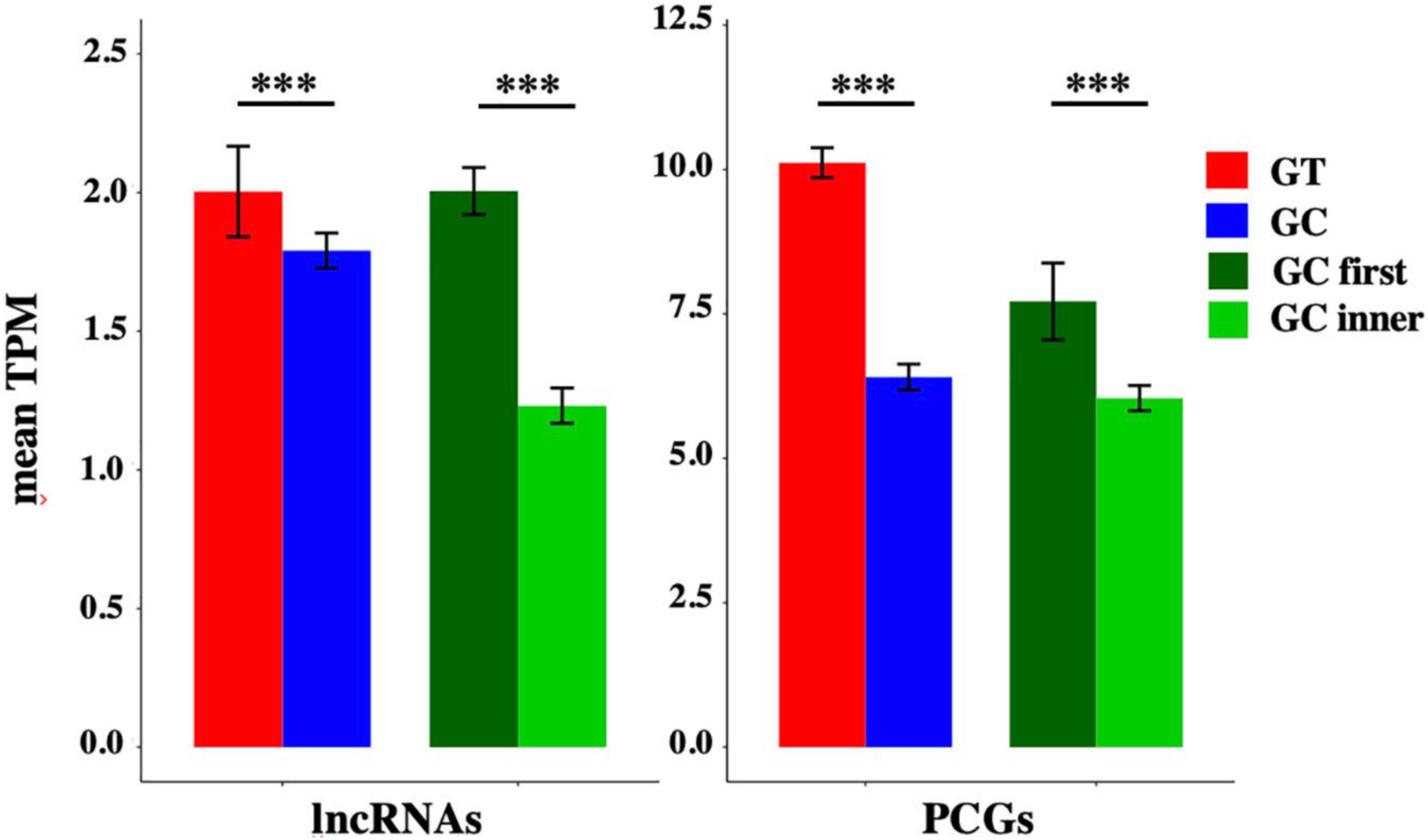
Expression of GC-AG- and GT-AG-containing transcripts. Bar graph representing the expression of lncRNAs and PCGs transcripts containing GC-AG- or GT-AG-introns and between transcripts containing a GC-AG intron in the first or inner position. The expression of transcripts was calculated as mean TPM combining expression data from 10 different tissues together. ***p < 0.001.

The mean TPM of transcripts containing a GC-AG intron in the first position appeared always higher with respect to transcripts having a GC-AG intron in inner positions, both in lncRNAs and PCGs. Considering the combination of all tissues, the mean TPM of GC-first introns lncRNAs resulted significantly higher with respect to GC-inner introns (lncRNAs: GC-first mean TPM 2.00 vs. GC-inner mean TPM 0.58, Wilcoxon test p-value < 2.2 × 10–16; PCGs: GC-first mean TPM 7.71 vs. GC-inner mean TPM 6.04, Wilcoxon test p-value = 5.4 × 10–11) (Figure 3). Interestingly, in some cases the expression levels of transcripts with the GC-AG intron located as the first resulted higher than GT-AG-containing transcripts especially in lncRNAs (i.e., in anterior cingulate cortex, amygdala, lung, skin, and spleen in lncRNAs and in heart for PCGs) (Supplementary Figure S4 and Supplementary Table S19).

These results suggest that the presence of a GC-AG intron may affect transcripts expression by reducing their overall transcription levels, both in lncRNAs and PCGs. Moreover, GC-AG introns may have a different effect on transcript expression levels depending on where they are located as transcripts harboring a GC-AG intron in their first intron showed overall higher expression levels with respect to transcripts with an inner GC-AG intron.

Nevertheless, these data must be taken with caution as: (i) the high variability in expression profiles, that is a common feature of both lncRNAs and PCGs, could affect mean TPM calculation especially for those categories containing a small number of transcripts, and (ii) transcripts containing a GC-AG intron often differ from GT-AG ones for other alternative splicing events which could possibly make differences in expression levels not univocally attributed to the presence of a GC intron.

### Multi-Species Conservation of GC-AG Introns

In human, GC-AG introns were present in 1224 lncRNAs and in 1934 PCGs, representing the 7.8 and 9.7% of each type of genes, respectively. In mouse, GC-AG introns were present in 473 lncRNAs and in 1669 PCGs, representing the 3.8 and 7.6% of each type of genes, respectively. The great majority of transcripts included one single GC-AG intron, especially for lncRNAs; few PCGs owned more than two GC-AG introns per transcript.

Based on the human-mouse ortholog information provided by the Ensembl project16, a total of 908 PCGs were conserved between the two species, thus accounting for a considerable fraction of total GC-AG containing genes (47% of human GC-AG containing genes; 54% of mouse GC-AG containing genes). Remarkably, in more than 75% of cases the GC-AG introns also shared the same ordinal position in the homologous genes.

Interestingly, we found many examples in which the conservation of the GC-AG introns together with their relative position inside the gene was not limited to mouse but it extended across evolutionary distant species. For example, the GC-AG splice sites of human ABI3BP (ABI family member 3 binding protein) and NDUFAF6 (NADH:ubiquinone oxidoreductase complex assembly factor 6) genes were shown to be conserved in chimp, macaque, mouse, rat, dog, cow, pig, chicken, fugu, and zebrafish (Figure 4). Moreover, the ordinal position of the GC-AG intron was also conserved: in the ABI3BP gene, GC-AG introns was always the first intron in all cases and in the NDUFAF6 gene, the GC-AG intron conserved its position in intron 6 in all species. The GC-AG splice sites of the human genes BLVRB (biliverdin reductase B) and AZI2 (5-azacytidine induced 2) were shown to be conserved in first and inner introns of mammals, respectively, while the canonical GT was found in chicken, fugu and zebrafish (Supplementary Figure S3). Despite the assessment of the conservation of lncRNAs was hindered by the lack of annotation in most species, a number of conserved GC-AG splice junctions between human and mouse was determined. Indeed, the TMEM51-AS1 (TMEM51 antisense RNA 1), the MALAT1 (metastasis associated lung adenocarcinoma transcript 1) and the NEAT1 (nuclear paraspeckle assembly transcript 1) genes contained a first GC-AG intron in both species whereas the JPX (JPX transcript, XIST activator) gene contained an inner GC-AG intron in both human and mouse.

**Figure 4.**
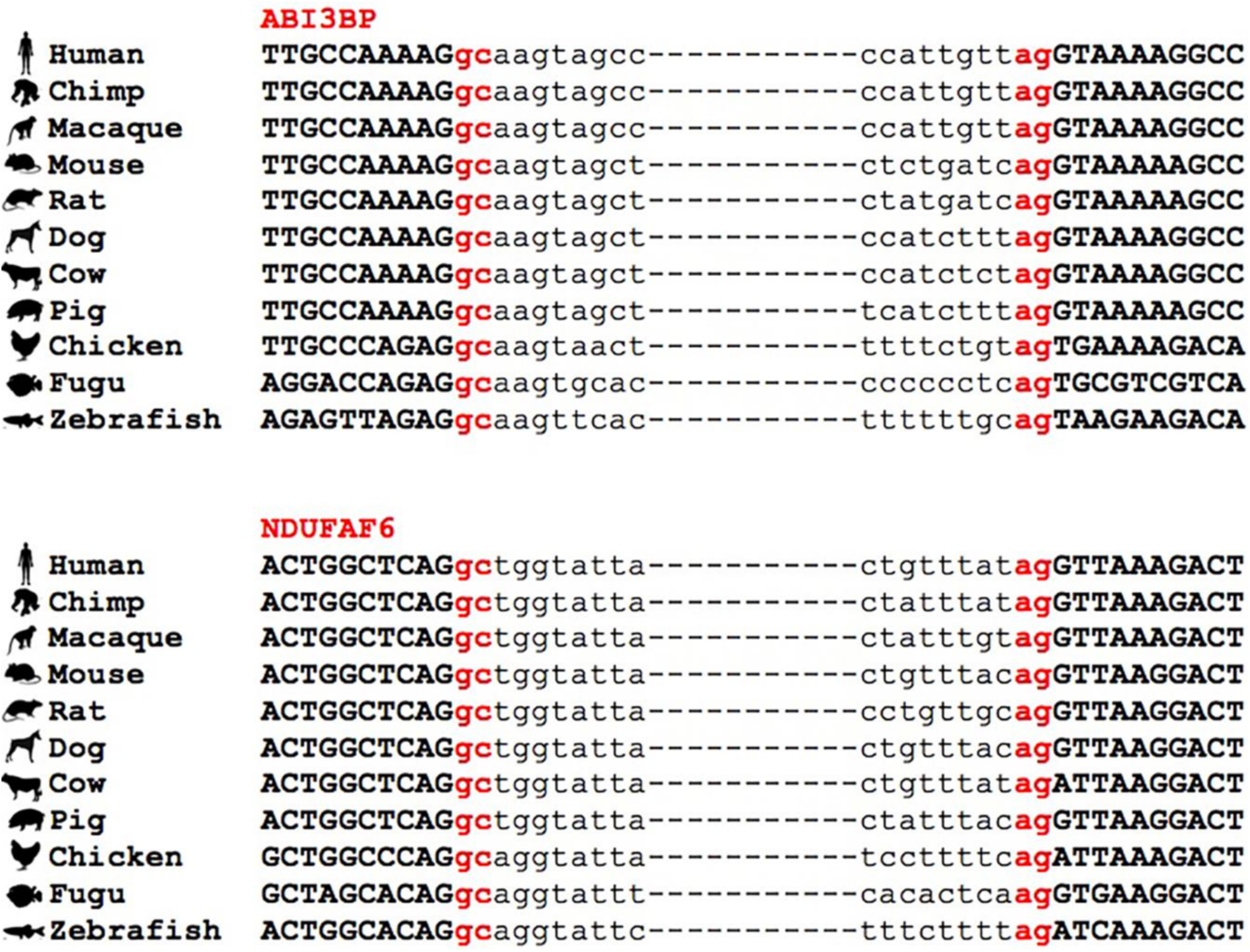
Conservation of GC-AG introns across multiple species. Multiple sequence alignment of GC-AG splice sites in the first intron of ABI3BP gene and the intron 6 of NDUFAF6 gene across the 11 species indicated.

The high conservation of the GC-AG introns between human and mouse and across multiple species could hint toward their functional importance and suggest their involvement in specific biological processes.

### Biological Processes Enrichment in GC-AG Containing Genes

In order to assess if the presence of a GC-AG intron may represent a regulatory motif involved in specific biological processes, we performed an enrichment analysis of Gene Ontology (GO) terms of human and mouse PCGs. By means of the DAVID Functional Annotation Tool (Huang et al., 2009) and the PANTHER Overrepresentation Test (Mi et al., 2019), we selected only those terms that resulted significantly enriched in both species and by both tools (Figure 5 and Supplementary Table S20).

**Figure 5.**
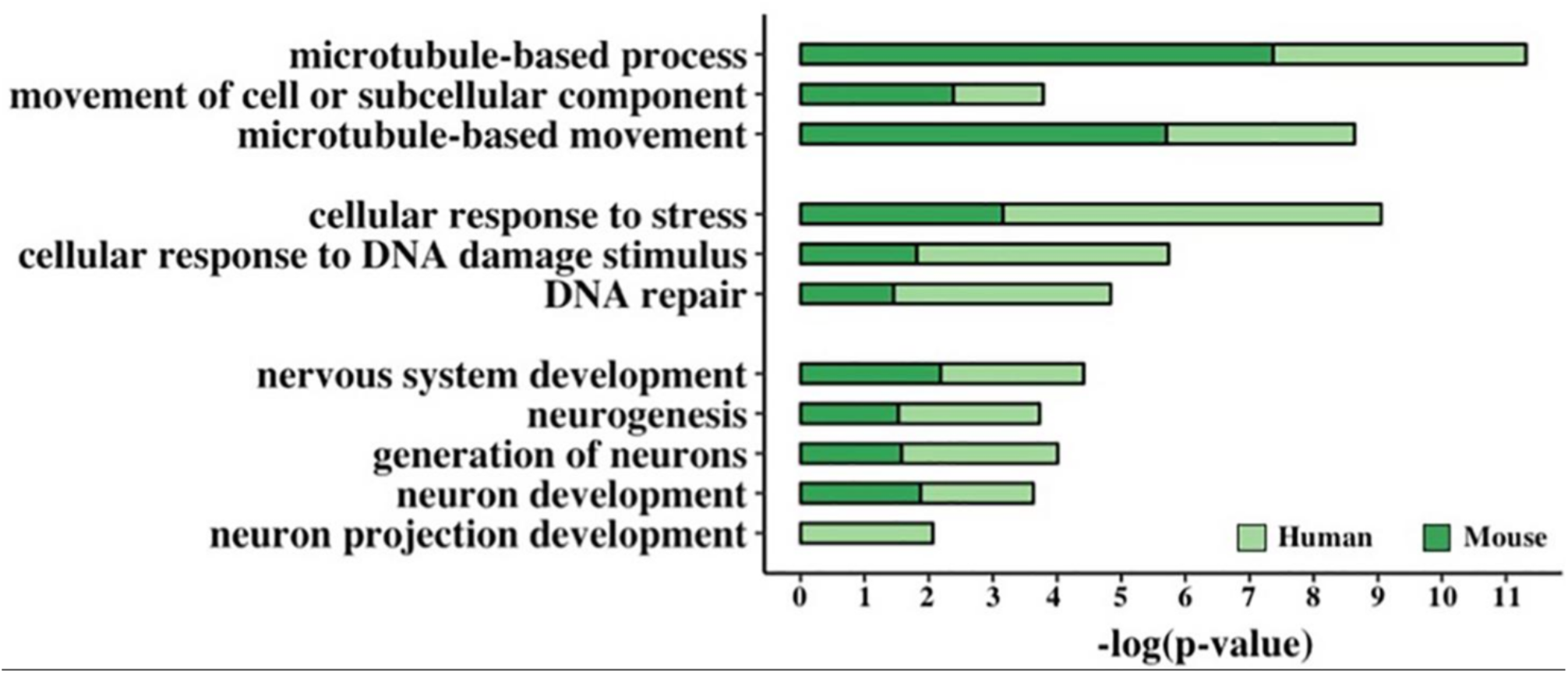
Functional enrichment analysis of GC-AG-containing genes. Bar graph representing the GO terms found significantly enriched in GC-AG containing PCGs. The GO term name is indicated on the Y-axis, and the (–)log10 of the p-values is indicated on the X-axis.

This resulted in the identification of three groups of related terms in the biological process ontology. The first group comprised the GO term “microtubule-based movement” and its ancestors “movement of cell or subcellular component” and “microtubule-based process” and included 221 human and 176 mouse genes. Despite very little is known about the biological processes in which lncRNAs are involved, at least two of the GC-AG-containing lncRNAs were described to have a role in the regulation of the movement of cells or subcellular components: the MEG3 (maternally expressed 3) gene (Wang et al., 2018; Xu et al., 2019) and the SOX2-OT (SOX2 overlapping transcript) gene (Wang et al., 2017). The second group contained the GO term “DNA Repair” and its ancestors “cellular response to DNA damage stimulus” and “cellular response to stress” and accounted for 257 human and 179 mouse genes. Interestingly, two of the GC-AG-containing lncRNAs were described to be involved in DNA repair: the MALAT1 gene (Hu et al., 2018) and the NEAT1 gene (Adriaens et al., 2016). In the third group, the GO term “neuron projection development” with its ancestors “neuron development,” “generation of neurons,” “neurogenesis,” and “nervous system development” were included and contained 273 and 220 human and mouse genes. Several lncRNAs with a GC-AG intron were described to play a role in neuron development and growth like the MEG3 gene (You and You, 2019), the NEAT1 gene (Barry et al., 2017), the SOX2-OT gene, the GDNF-AS1 (GDNF antisense RNA 1) gene and the MIAT (myocardial infarction associated transcript) (Clark and Blackshaw, 2014). All the reported GO terms resulted significantly enriched after correction for multiple testing (Figure 5 and Supplementary Table S20).

## DISCUSSION

In this study, we report a genome-wide comparison of genomic and splicing features of long non-coding and protein-coding genes in human and mouse. Being based on GENCODE releases 27 and M16, our analysis considered a conspicuously higher number of genes with respect to previous studies (Cabili et al., 2011; Derrien et al., 2012) and it was strengthened by the comparison between the two species.

The characterization of the genomic features revealed differences in the genetic architecture between long non-coding and PCGs in both human and mouse. We found that lncRNAs were shorter than protein-coding ones in both species in agreement with previous studies (Ravasi et al., 2005; Cabili et al., 2011; Derrien et al., 2012); however, this was not only due to the lower number of exons but also to the shorter length of exons in lncRNAs. The shorter length and the limited number of exons in lncRNA genes might be attributed to their incomplete annotation as their low expression level and high tissue specificity hampers the complete characterization, as suggested by the studies of Lagarde et al. (2016, 2017). Nevertheless, our results did not appear to be driven by this bias as we used a recent and more complete GENCODE release, whose annotation was based on stronger experimental and computational evidence (Frankish et al., 2019) and they were confirmed in six more lncRNAs annotation datasets (FANTOM5, NONCODEv5, BIGTranscriptome, MiTranscriptome, LNCipedia, and LncBook). In particular, the FANTOM CAT robust lncRNA annotations specifically providing accurate annotations of transcripts’ TSS and 5′ ends through the Cap Analyses of Gene Expression (CAGE) method, and the BIGTranscriptome dataset employing both CAGE and poly(A)-position profiling by sequencing (3P-seq) to assess 5′ and 3′ end completeness, indicated that our results are not subjected to the bias of incompleteness. It is nevertheless interesting to note that the reduction in size in lncRNAs affect those portions of the gene mainly involved in gene expression regulation. The length of the first exon was described to be related to transcription efficiency and it was reported that short first exons could promote transcriptional accuracy as they exhibit a more concentrated assembly of transcription factors near transcription start sites (Bieberstein et al., 2012). Moreover, last exons tend to be longer than first and inner exons due to the presence of 3′UTR sequences, essential for the regulation of multiple aspects including nuclear export, cytoplasmic localization, stability, and translational efficiency (Kalari et al., 2006; Andreassi and Riccio, 2009). Thus, our results hints toward a difference in the regulatory potential contained in the first and last exons of lncRNAs. Taken together, our data suggested that the difference in gene architecture between lncRNAs and PCGs could imply their involvement in different mechanisms of genomic control and gene expression regulation.

The characterization of splicing features revealed a significant enrichment of introns harboring GC-AG splice sites in lncRNAs of both species. GC-AG splice sites were generally considered as a non-canonical variant of the major U2-type GT-AG splice junctions, accounting for 0.865 and 0.817% in human and mouse genomes, respectively (Sheth et al., 2006; Parada et al., 2014). In agreement with what was previously reported, we assessed the same frequency of GC-AG introns in both species when considering only PCGs (0.83% in human and 0.81% in mouse). When lncRNAs were taken into account, the frequency of GC-AG splice sites resulted more than three time higher in human and more than two times higher in mouse, accounting for 3.0 and 1.9% of their total splice junctions. Notably, the enrichment of GC-AG splice sites did not appear to be evenly distributed, as it emerged more prominent in the first intron of both types of genes. In human, GC-AG first introns corresponded to 4.2 and 1.2% of total first introns of lncRNAs and PCGs, respectively. The same trend was observed in mouse in which a higher ratio of GC-AG splice junctions were found in the first intron in both lncRNAs (2.4%) and PCGs (1.2%). The enrichment of GC-AG introns in lncRNAs and their preferential position in the first intron did not appear to be driven by a mis-annotation bias as the same trend was also observed in the FANTOM5 dataset. The same enrichment was also assessed in the lower organisms D. melanogaster and C. elegans, despite this analysis could not be conclusive due to incomplete annotations and limited number. The significant increase of GC-AG introns in lncRNAs, together with their non-random distribution along the gene, led us to hypothesize that they may represent unique regulatory elements. The preferential localization of GC-AG splice sites in the first intron provided a clear indication of their role in gene expression regulation. Indeed, first introns were described to possess particular regulatory features, as they were shown to be more conserved with respect to inner introns and to be enriched in epigenetics marks associated with active transcription, such as H3K4me3 and H3K9ac (Bieberstein et al., 2012; Park et al., 2014), thus being likely involved in gene expression and splicing regulation. In many cases, first introns were demonstrated to be responsible for transcription initiation and increase of mRNA transcriptional rates (Rose, 2019). Moreover, the binding of the U1-complex to 5′ss was demonstrated to be involved not only in splicing regulation but also in polyadenylation control and in regulation of gene expression through its interaction with promoter (Berg et al., 2012; Almada et al., 2013; Singh and Singh, 2019) suggesting that the non-canonical GC 5′ss could in some way perturb this mechanism of action.

GC-AG introns displayed distinctive splicing features in comparison with GT-AG introns, in particular when located in the first intron of lncRNAs. Introns harboring GC-AG splice sites appeared significantly shorter than GT-AG introns, in both lncRNA and PCGs. This trend was more prominent in human GC-AG first introns, having an average length of ~6.7 kb in lnc-genes and ~9 kb in pc-genes, and being significantly shorter than GT-AG first introns (~13 and ~15 kb in lncRNAs and PCGs genes, respectively). In addition to their shorter length, GC-AG splice sites appeared significantly weaker than GT-AG ones. A reduction in the 5′ss strength of GC-AG introns was expected because of the mismatch at position +2 with the U1 snRNA consensus. Nevertheless, the reduction of 5′ss strength was more evident in GC splice sites of lncRNAs rather than in PCGs and it was more prominent in the first intron rather than in inner ones. Similar results were obtained for 3′ss, whose average weight-matrix scores for GC-AG introns appeared significantly lower compared to GT-AG junction, especially when located in lncRNAs first introns. Interestingly, the Spearman correlation test demonstrated a positive correlation among intron length and 5′/3′ss strength for the first intron of lncRNAs, thus implying the enrichment of short and very weak first introns in this class of molecules.

It was suggested that the base pairing between 5′ss and U1 regulates alternative versus constitutive splicing, hence suggesting that weak splice sites are more prone to undergo alternative splicing (Stamm et al., 1994; Sorek et al., 2004). In agreement with previously reported data (Kralovicova et al., 2011), our analysis at the gene level confirmed that GC-AG containing genes were more prone to alternative splicing than genes harboring GT-AG introns. In addition, our analysis suggested that GC-AG introns might be preferentially involved in alternative 5′ss splicing and alternative polyadenylation events, thus indicating a specific role that will require further investigations. Churbanov et al. (2008) demonstrated that an excess of GT to GC 5′ss conversions occurred both in primates and rodents, hypothesizing that the accumulation of GC sites in mammals might arise from positive selection favoring alternative splicing. Moreover, GC-AG introns were found to be strongly overrepresented in recent intron gain events occurring in segments associated with repetitive sequences that are highly alternatively spliced (Zhuo et al., 2007). Taken together, these results further supported the role of GC-AG introns as regulatory elements putatively involved in the control of alternative splicing events. How GC-AG introns could contribute to increase alternative splicing levels and polyadenylation regulation will require further investigations.

A preliminary analysis of RNA-seq data of 10 different human tissues from the GTEx project, allowed us to highlight a putative effect of GC-AG introns on gene expression profiles. Indeed, the overall expression of GC-AG introns containing transcripts appeared lower with respect to GT-AG ones thus suggesting they may have a reduction effect on gene expression. More interestingly, our data suggested that GC-AG introns located as first behave differently as they demonstrated higher level of expression with respect of transcripts containing an inner GC-AG intron thus underlining their peculiar regulatory role depending on the position. Despite we are aware that these data must be taken with caution as they may be biased in many ways, they represent a first experimental evidence of the effect of GC-AG introns at gene expression level.

Despite the percentage of GC 5′ss is relatively small, the number of genes containing at least one GC-AG intron is not irrelevant, as they account for about 10% of pc-genes and 8% of lncRNAs in human (in mouse: about 8% of PCGs and 4% of lncRNAs). The relevance of GC-AG-containing genes emerged also from the analysis of their conservation: about 50% of GC-AG containing PCGs resulted conserved between human and mouse which could also be related to late intron gain events. Furthermore, in the majority of conserved PCGs (75%), the ordinal position of GC-AG introns was also conserved. As 25% of GC-AG introns do not have the same ordinal position, this could also argue about recent intron gain occurrence. Moreover, in many instances the GC-AG splice sites appeared to be conserved not only in the mouse genome but also in other species and across large evolutionary distance. The evaluation of the conservation of GC-AG splice sites in lncRNA genes was hindered by their current incomplete annotation in many species. However, among the well-studied and annotated lncRNAs, we still could identify examples of the conservation of GC-AG splice sites between human and mouse. Indeed, the two well characterized nuclear lncRNAs NEAT1 and MALAT1 juxtaposed on human chromosome 11 (on chromosome 19 in mouse) share similar gene features: both are transcribed in long unspliced isoforms as well as in shorter and spliced transcripts starting from the same promoter. Moreover, both NEAT1 and MALAT1 shorter transcripts contain a GC-AG first intron in human and mouse, thus suggesting similar regulatory functions.

The functional enrichment analysis of human and mouse PCGs provided further evidence that GC-AG introns could represent a specific regulatory motif as it revealed a significant enrichment of GO terms related to DNA repair, neurogenesis, and microtubule-based movements. Despite the enrichment analysis for lncRNA genes was obstructed by the lack of their functional annotation, we reported several examples of the involvement of lncRNA genes harboring a GC-AG introns in these biological processes. This analysis suggested that GC-AG introns may be involved in the expression control of genes involved in specific cellular functions, reasonably needing a concerted regulation.

In few cases, the functional relevance of GC-AG introns was already demonstrated. In the study of Farrer et al. (2002) it was demonstrated that the weak GC 5′ss located in intron 10 of the Collagen alpha-2(IV) chain (let-2) gene in C. elegans was essential for developmentally regulated alternative splicing, and that its replacement with a stronger GT splice site suppressed the alternative splicing regulation occurring during embryos development. In the inhibitor of growth family member 4 (ING4) gene, the selection between a weak GC 5′ss or a near-located canonical GT was shown to result into alternative transcript isoforms which diverged for the presence of a nuclear localization signal thus affecting the subcellular localization of the encoded protein (Tsai et al., 2008). In the work of Palaniswamy et al. (2010), a single nucleotide polymorphism converting a 5′ss GT to GC, present with varying frequencies in different mouse strains, was shown to be responsible for an alternative splicing event affecting the length and the translational efficiency of the GLI-Kruppel family member GLI1 (Gli1) gene in mouse. Moreover, for the PR/SET domain (PRDM) gene family in human (Fumasoni et al., 2007) and for the starch synthase (SS) gene family in rice (Chen et al., 2017) the activation of a GC 5′ss was shown to contribute to the diversification and the evolution of both gene families.

It is today clear that organisms complexity does not correlate with genome size or gene content, but it is instead more consistently related to the level of gene expression regulation. An higher level of gene regulation is thought to ensure the development of more sophisticated capabilities of higher organisms, despite the fact that the number of PCGs is similar in evolutionary distant species. Furthermore, the amount of alternative splicing, which allows the production of a wide variety of proteins starting from a smaller number of genes, is known to be positively correlated with eukaryotic complexity (Bush et al., 2017; Schaefke et al., 2018). Moreover, the amount of transcribed ncDNA resulting in the production of a large collection of ncRNAs mainly involved in the regulation of gene expression, is known to increase together with organisms complexity (Liu et al., 2013; Jandura and Krause, 2017). As it occurs for alternative splicing and for non-coding transcripts, also the frequency of GC-AG splice sites was reported to correlate with metazoan complexity (Sheth et al., 2006), hence supporting the idea that this class of introns may represent a new layer of gene regulation. Interestingly, the conversion of donor splice sites from GT to GC was demonstrated to be an evolutionary driven mechanism, putatively due to the increased number of alternative splicing events occurring at weak GC-AG introns (Abril et al., 2005; Churbanov et al., 2008).

Taken together, our data suggested that GC-AG introns represent new regulatory elements mainly associated with lncRNAs and preferentially located in their first intron. Their increased frequency in higher organisms suggested that they could contribute to the evolution of complexity, adding a new layer in gene expression regulation. How they exerted their regulatory role remains to be further investigated despite preliminary evidence suggested that they could favor alternative splicing. The elucidation of the mechanisms of action of GC-AG introns could contribute to a deeper and better understanding of gene expression regulation and could address the comprehension of the pathological effects of mutations affecting GC donor sites contained in several disease-causing genes.

## Acknowledgments

We thank Prof. C. Calvio, Dr. S. Sabbioneda, Dr. R. Alfieri, and Dr. C. Mondello for critical reading of the manuscript. MA and LC are enrolled in the Ph.D. program in “Genetics, Molecular and Cellular Biology” of the University of Pavia, Italy.

