## Additional file 1 for "GC-AG Introns Features in Long Non-coding and Protein-Coding Genes Suggest Their Role in Gene Expression Regulation"

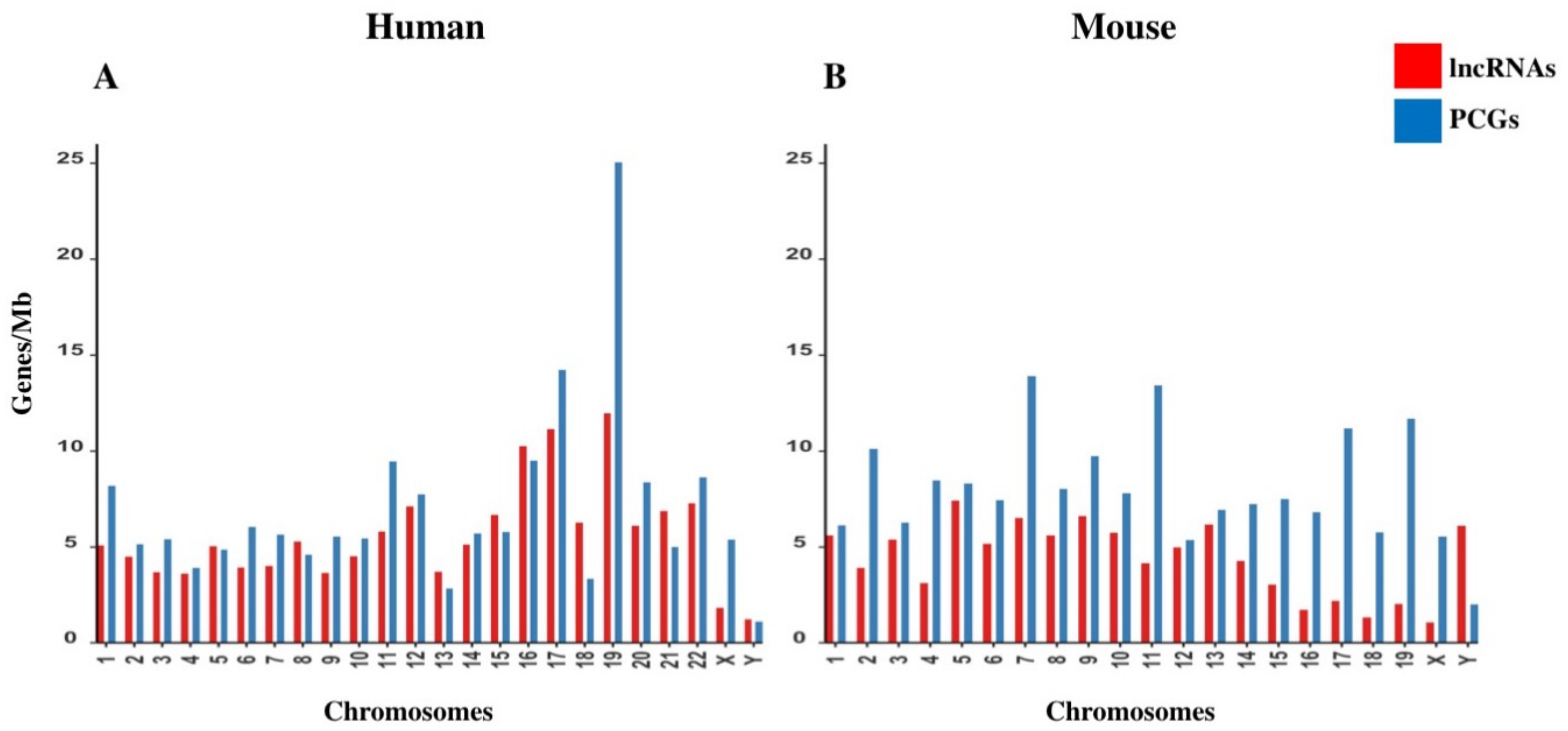


**Supplementary Figure 1 -** Gene densities of lncRNAs and PCGs in human (A) and mouse (B) across chromosomes. Densities were reported as number of genes per Megabase (Mb).

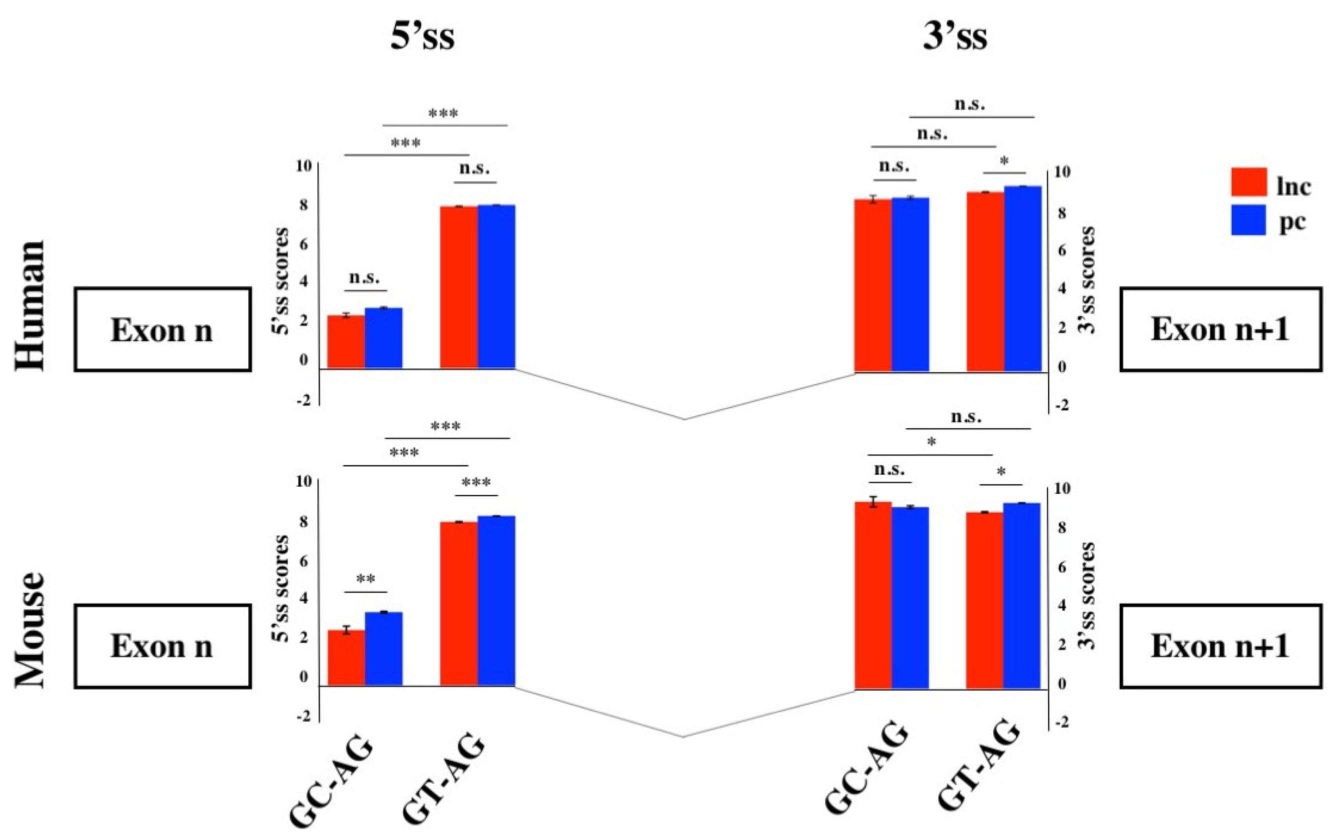


**Supplementary Figure 2 - Splice junctions strengths of inner introns.**
Schematic representation of the average scores of 5' and 3'ss strengths (weight matrix scores) of lncRNAs and PCGs in human and mouse for GC-AG and GT-AG inner introns.

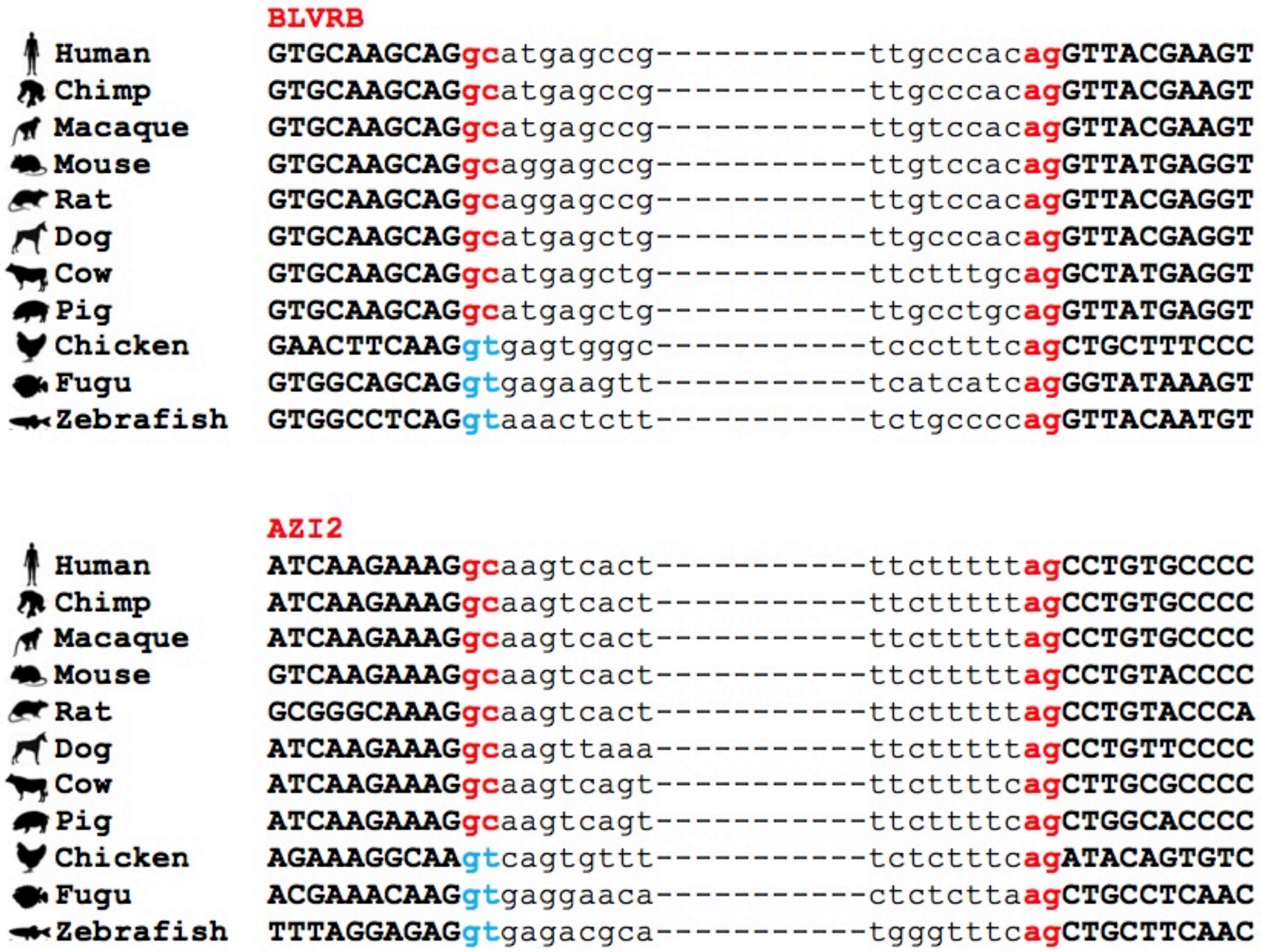


**Supplementary Figure 3 - Conservation of GC-AG introns across multiple species**. Multiple sequence alignment of GC-AG splice sites in the first intron of *BLVRB* gene and the inner intron of *AZI2* gene across 11 species.

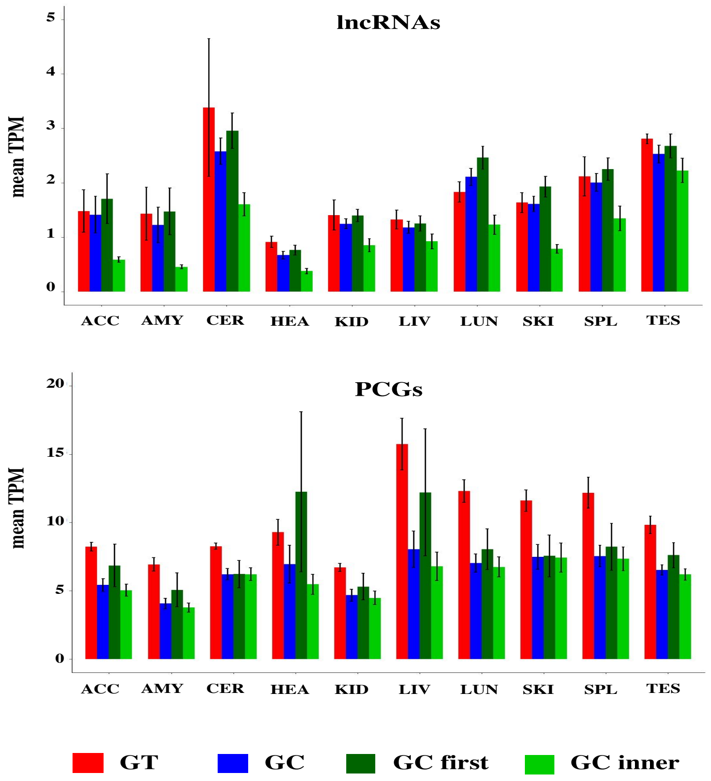


**Figure 4 – Expression of GC-AG- and GT-AG-containing transcripts.** Bar graph representing the expression of lncRNAs and PCGs transcripts in different human tissues (acc: anterior cingulate cortex; amy: amygdala; cer: cerebellum; hea: heart; kid: kidney; liv: liver; lun: lung; ski: skin; spl: spleen; tes: testis)Transcripts were divided as containing GC-AG- or GT-AG-introns and between transcripts containing a GC-AG intron in the first or inner position. The expression of transcripts was calculated as mean TPM combining expression data from 10 different tissues together.
